# Evaluating scientific theories as predictive models in language neuroscience

**DOI:** 10.1101/2025.08.12.669958

**Authors:** Chandan Singh, Richard J. Antonello, Sihang Guo, Gavin Mischler, Jianfeng Gao, Nima Mesgarani, Alexander G. Huth

## Abstract

Modern data-driven encoding models are highly effective at predicting brain responses to language stimuli. However, these models struggle to *explain* the underlying phenomena, i.e. what features of the stimulus drive the response? We present <u>Q</u>uestion <u>A</u>nswering encoding models, a method for converting qualitative theories of language selectivity into highly accurate, interpretable models of brain responses. QA encoding models annotate a language stimulus by using a large language model to answer yes-no questions corresponding to qualitative theories. A compact QA encoding model that uses only 35 questions outperforms existing baselines at predicting brain responses in both fMRI and ECoG data. The model weights also provide easily interpretable maps of language selectivity across cortex; these maps show quantitative agreement with meta-analyses of the existing literature and selectivity maps identified in a follow-up fMRI experiment. These results demonstrate that LLMs can bridge the widening gap between qualitative scientific theories and data-driven models.

## 1 Introduction

The advent of powerful deep learning models has provided neuroscientists with new tools for modeling how the brain encodes information across language, vision, and audition (*1, 2, 3*). One such tool is the *language encoding model*, which allows neuroscientists to predict the relationship between a language stimulus and its evoked brain response with unprecedented accuracy (*4,5,6,7*). However, these increasingly large, black-box models are unable to succinctly *explain* their predictions. This has challenged our notion of what it means to understand a natural phenomenon: if our model of the brain is as inscrutable as the brain itself, does that model advance science? Or must our understanding take the form of a scientific theory that can be expressed in words?

To meet this challenge, we present <u>Q</u>uestion <u>A</u>nswering encoding models, a method for evaluating qualitative scientific theories as predictive models. First, theories about language in the brain (e.g. *Some brain areas selectively respond to language about time*) are recast as yes/no questions about a piece of text (e.g. *Does the input mention time?*). An LLM is then used to annotate natural language stimuli from a neuroimaging experiment by answering these questions about each segment of text. We then fit linear models that use these annotations as features to predict brain responses, yielding interpretable, highly predictive encoding models. Using as few as 35 questions, QA encoding models outperformed even state-of-the-art black-box encoding models for both fMRI and ECoG.

Each feature in a QA encoding model corresponds to a single theory. Thus each voxel’s linear model weight for a particular feature is highly interpretable, showing which brain areas’ responses match the corresponding theory. We confirmed the reliability of these maps by showing that they match meta-analyses of the existing literature (*8*). We further evaluated these maps using follow-up fMRI experiments. Specifically, we used generative causal testing (*9*), which constructs synthetic stories where each paragraph corresponds to a particular theory. We then measured the average fMRI response to each paragraph and found strong agreement with the QA maps. Our results demonstrate the power of QA encoding models to quantitatively evaluate qualitative scientific theories using large-scale data.

## 2 QA encoding models for fMRI are highly accurate and compact

To construct QA encoding models, we began by enumerating a set of 606 plausible neurolinguistic theories expressed as yes/no questions. We generated these questions by first manually writing many examples and then prompting GPT-4 (*10*) to create more using various strategies, like having GPT-4 list semantic properties of narrative sentences, summarize n-grams from the training data, or generate questions similar to single-voxel explanations found in a prior work (*11*) (Fig. 1a). We then used an ensemble of LLMs to annotate each 10-gram in 20 hours of narrative stories with the answer to each question (Fig. 1b, see Methods). These answers serve as an interpretable binary-valued text embedding (*12*). Next, these embeddings were used to fit a linear encoding model for each voxel in 3 human subjects from passive language listening fMRI data (*13*). The encoding models were then tested by predicting held-out fMRI data and computing the correlation between predictions and actual data (Fig. 1c).

**Figure 1:**
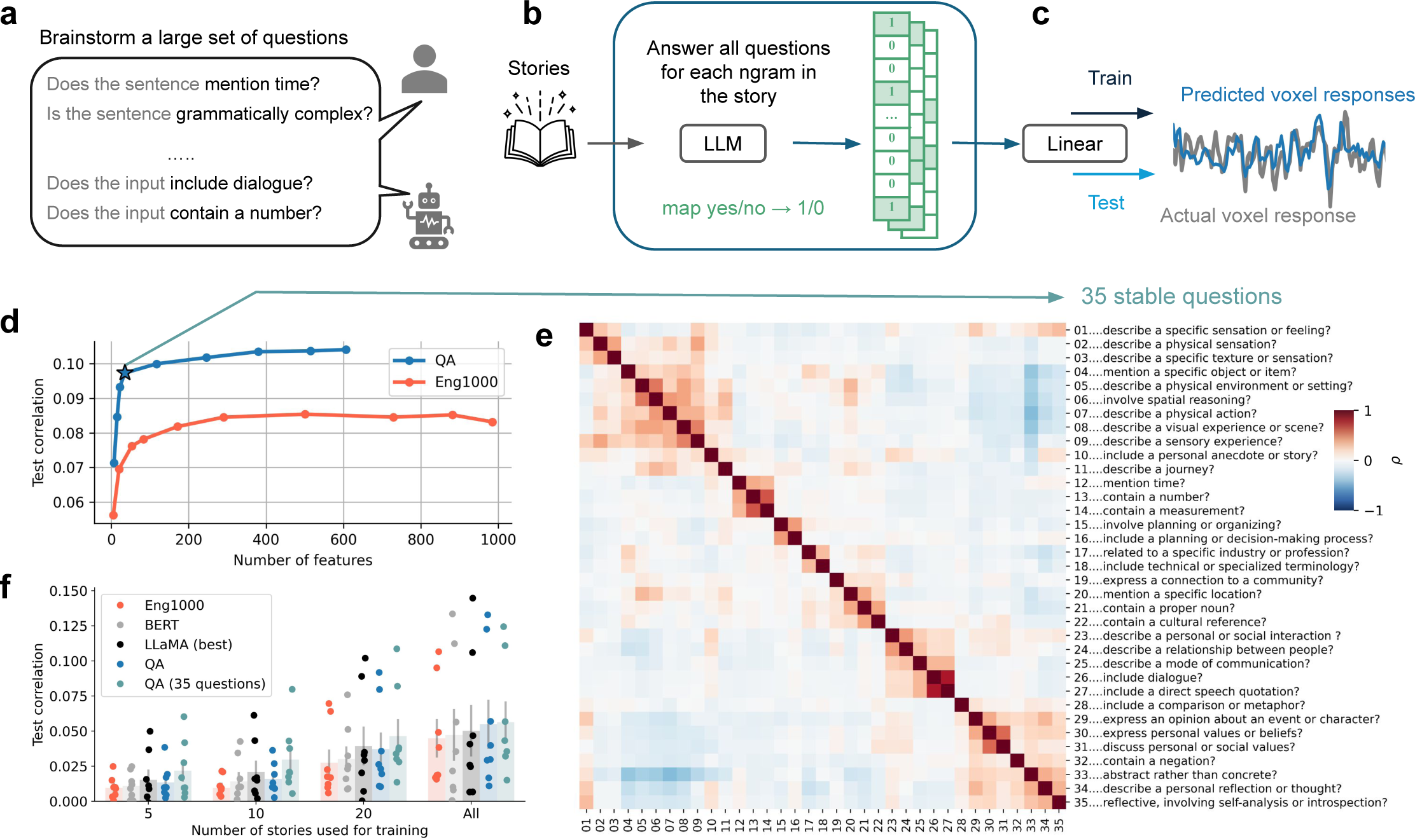
QA encoding models effectively predict fMRI voxel responses from LLM answers to questions. (a) We first enumerated a set of 606 qualitative theories cast as yes/no questions through brainstorming and GPT-4 assistance. (b) 20 hours of narrative stories were converted to 606 interpretable, binary features by having LLMs answer each question for every 10-gram in the stories. (c) These features were then used to build voxelwise encoding models. Voxelwise BOLD responses were recorded using fMRI as human subjects listened to the narrative stories. Regularized linear encoding models were fit to predict each voxel response from the binary features; encoding models were tested by predicting responses on held-out fMRI data. (d) To compress the resulting 606-question encoding models, sparse stability selection was used during fitting. Very few questions were required to achieve strong prediction performance: with only 35 questions, the QA model achieved a test correlation of 0.097, a 13.9% improvement over the best performance (averaged over 3 subjects) attained by the much larger Eng1000 baseline word embedding model. (e) The 35 selected questions and correlations between their answers on the 20 hours of stimuli. They span a variety of topics. (f) Average test encoding performance across cortex for the QA model and baselines on the three original subjects (20 hours of fMRI data each) and 5 additional subjects (5 hours each). The 35-question QA model outperformed the state-of-the-art black-box model (which uses hidden representations from the LLaMA family of LLMs) by 12.0% when trained on all the story data. The model’s compactness yields greater relative improvements in data-limited scenarios; when trained on only 5 stories per subject it outperforms the baseline LLaMA model by 43.3%.

To evaluate which of the 606 questions were useful for predicting brain responses, we employed sparse stability selection (*14*) of regressors during model fitting (Fig. 1d). This process selected questions that were individually strong predictors (Fig. S1a) and that represented a diverse set of information (Fig. S1b). The QA model that selected only 35 questions (hereafter QA-35) achieved a test correlation of 0.097, already a 13.9% improvement over the best performance attained by Eng1000, a baseline interpretable word embedding model consisting of 985 word-level features (*15*). We visualized the questions in QA-35 along with the correlation matrix of their answers across the stimulus and found that they span a variety of semantic concepts (Fig. 1e). A majority of the questions are loosely clustered into a handful of high-level categories: questions about *tactile sensations*, *visuospatial information*, *numerical information*, *planning*, *communication*, and *abstract beliefs or values*. As these categories are almost entirely what our model relies upon to make its inferences, they provide additional insight into the large-scale organization principles, such as grounded cognition (*16*), that govern the structure of cortical language representations (*15*).

We further evaluated the performance of QA-35 on held-out data for the original 3 subjects as well as 5 additional subjects that were not involved in the question selection. We found that it outperformed even black-box encoding models that use the hidden representations of an advanced LLM, LLaMA (*17*), by an average of 12.0% when models were trained on all available fMRI data (*7*) (Fig. 1f). The compactness of QA-35 additionally enables improved data efficiency, leading to larger relative improvements for scarce or low-quality fMRI training data. For example, when models were trained with only 5 stories (roughly 75 minutes of data), the relative improvement over the best baseline was 43.4%.

## 3 Interpreting cortical maps from QA encoding models

The compactness and linearity of QA-35 provide an opportunity to evaluate which areas of cortex are best captured by each qualitative theory. QA-35 predicts the BOLD response timecourse of each voxel as a weighted combination of the answers to 35 questions. Thus, if responses in some voxel are explained by a particular theory, the weight for the corresponding question in that voxel will be high. For example, a voxel that is selective for language about locations should have a high regression weight for the question

### Does the input mention a specific location?

Fig. 2a shows the encoding weights across the 35 questions for the left hemisphere of subject S02. The weights show many patterns that are consistent with known neuroscientific principles and theories. For the *locations* question (question 6), the map shows high weights for regions known to have selectivity for place concepts: retrosplenial cortex (RSC), the parahippocampal place area (PPA), and the occipital place area (OPA) (*18*). Visuospatial information strongly predicts activations in the occipitoparietal junction (*19*), whereas commmunication and speech information heavily predicts activations in inferior temporal (IT) lobe (*9*). Information about tactile sensations uniquely activates the supramarginal gyrus, which is home to somatosensory association cortex (*8*). See full weights for all 35 questions in Fig. S7.

**Figure 2:**
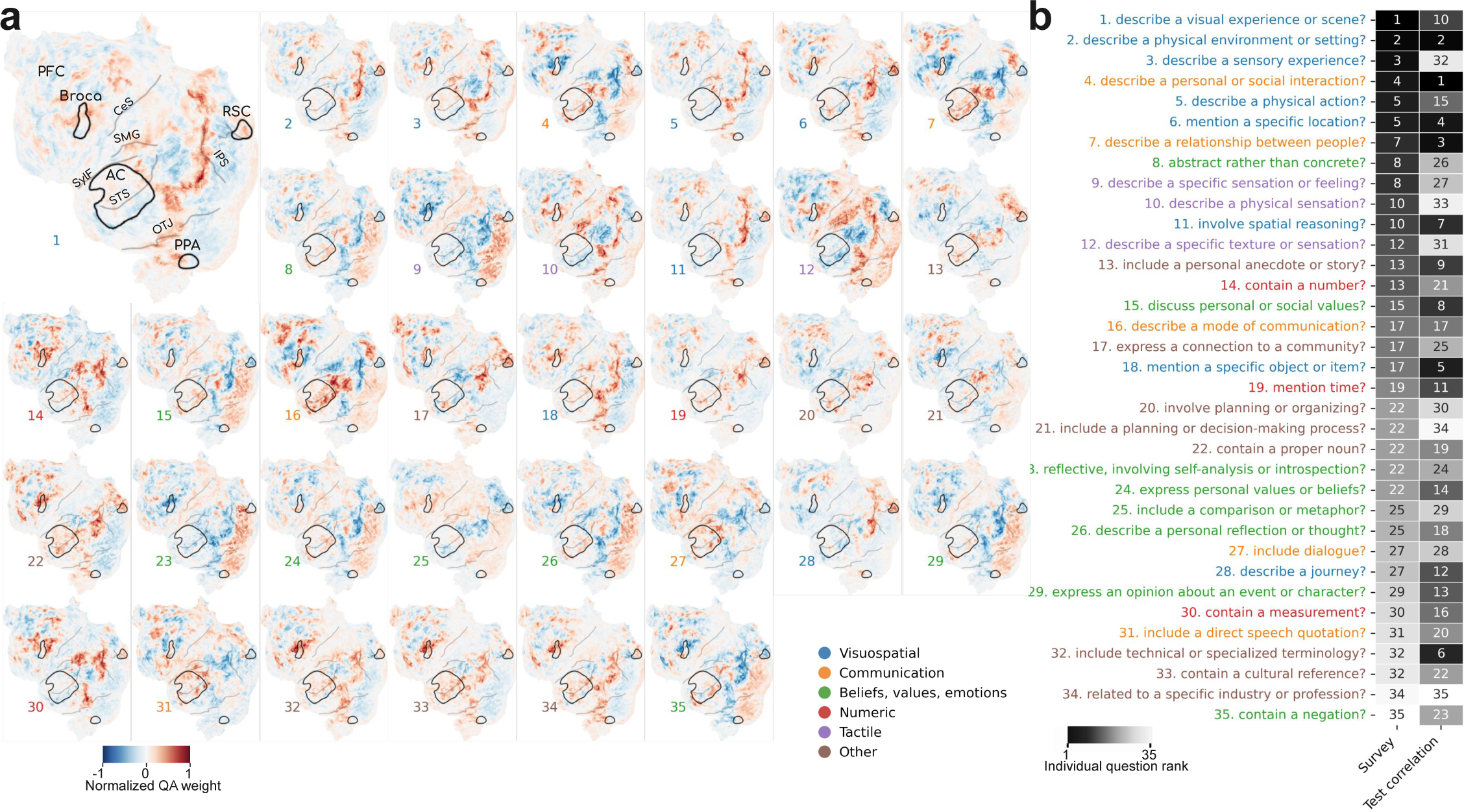
Interpreting cortical maps from QA encoding models. The 35-question QA encoding model in Fig. 1 yields weights that predict each voxel’s responses from each question’s answers; we can interpret these weights as cortical maps of semantic selectivity. (a) These semantic selectivity maps recover known selectivity patterns as well as uncovering new ones. For example, the weights for a question asking about whether the input mentions *locations* (question 6) have high values for regions known to have selectivity for place concepts: retrosplenial cortex (RSC), the parahippocampal place area (PPA), and the occipital place area (OPA). For brevity we show only the left hemisphere in a single subject (S02); see the full-cortex weights for multiple subjects in Section S3.3. Regions annotated are prefrontal cortex (PFC), Broca’s area, central sulcus (CeS), supramarginal gyrus (SMG), the Sylvian fissure (SylF), auditory cortex (AC), superior temporal sulcus (STS), occipitotemporal junction (OTJ), intraparietal sulcus (IPS), retrosplenial cortex (RSC), and parahippocampal place area (PPA). (b) To compare the findings in these selectivity maps against expert opinion, we collected the judgements of the importance of each of the 35 questions in QA-35 for predicting brain responses to language via an anonymous survey. We compared the average ranking of each question based on the survey (1 being the most important) to the ranking of each question based on its individual prediction performance, measured via test correlation. We found a somewhat positive correlation of 0.31 between the two rankings (𝑝 = 0.071). Many of the top questions ranked by experts were related to visuospatial concepts. Some questions, such as question 32 about *technical or specialized terminology*, are considerably more important for prediction than expert rankings suggested.

To compare the findings in these 35 selectivity maps against expert opinion, we conducted an anonymous survey asking researchers to provide their judgement of each question’s importance for predicting brain responses to language using a five-point Likert scale (e.g. 1 = “Not at all important”, 5 = “Extremely important”). The survey was sent out to four relevant mailing lists and yielded 12 responses; see full survey details in the Methods. Fig. 2b compares the average survey rating of each question (1 being the most important) to the question’s rank based on its individual prediction performance, measured via test correlation. We found a somewhat significant positive correlation of 0.31 between the two rankings (𝑝 = 0.071). Many of the top questions ranked by experts were related to visuospatial concepts. A handful of questions, such as *Does the input include technical or specialized terminology?* and *Does the input contain a cultural reference?* were considerably more important for prediction than expected by experts, likely because they are highly abstract and do not readily correspond to well-known theories about the drivers of cortical organization. These questions tended to correspond to complex patterns of activation in frontal lobe, near Broca’s area, as well as along the superior temporal sulcus. See full survey results in Fig. S6.

## 4 Quantitatively evaluating cortical maps from QA encoding models

To quantitatively evaluate the interpretation of QA-35 weights as cortical selectivity maps, we performed three robustness analyses. As a running example, we show the weights for the question *Does the input mention a specific location?*; see weights for subject S02 in Fig. 3a.

**Figure 3:**
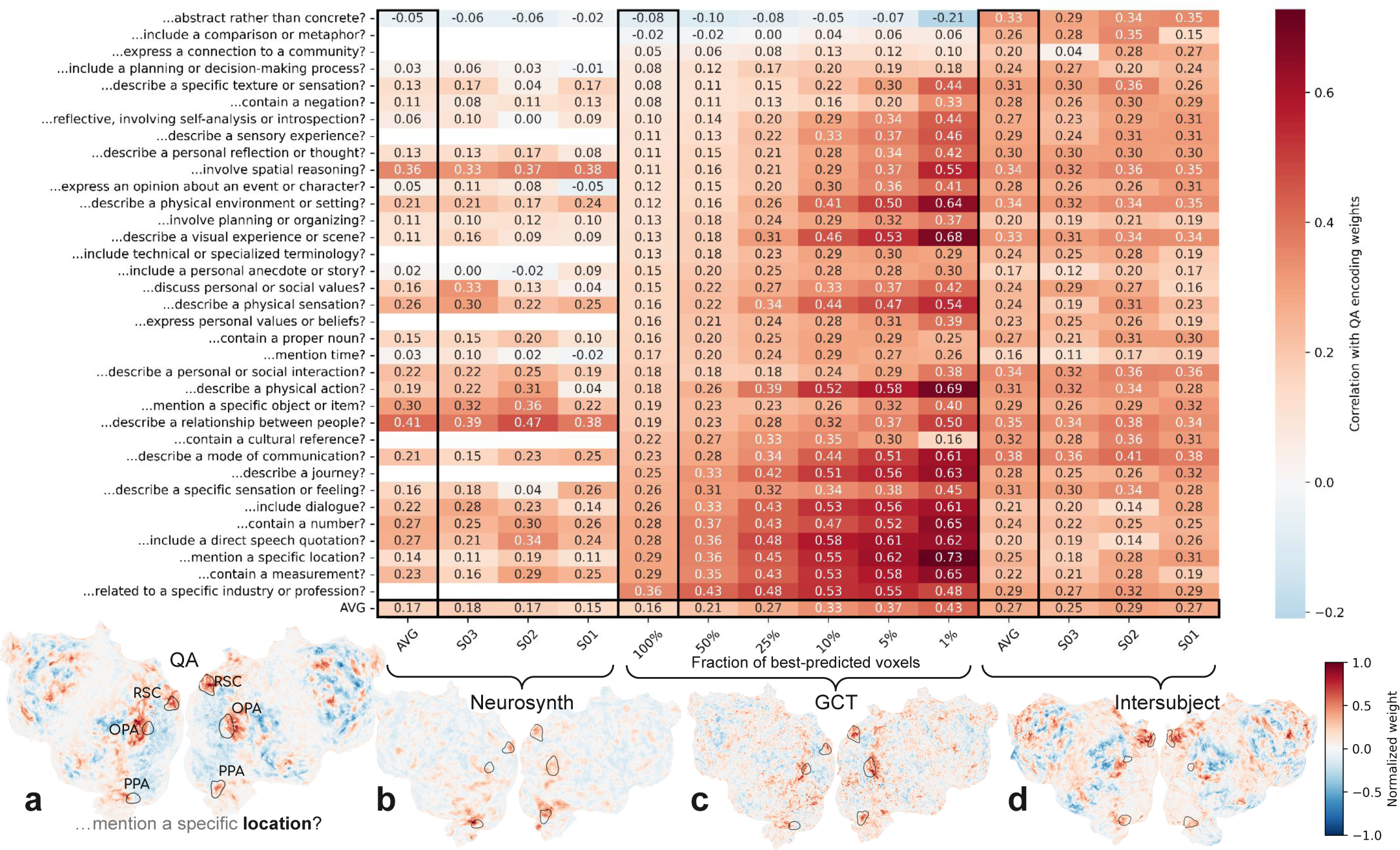
Quantitatively evaluating QA cortical selectivity maps corresponding to 35 stable questions. We evaluate the weights from the 35-question QA encoding model in Fig. 1 as cortical maps of semantic selectivity. (a) As a running example, we show the weights for a question asking about *locations* in subject S02 that displays selectivity for well-studied location-selective regions, such as RSC, OPA, and PPA. (b) To quantitatively evaluate whether the QA weights match known findings in the literature, we compared the QA weights against cortical maps from Neurosynth, a meta-analysis of the neuroscience literature that computed selectivity maps by testing the voxel-level association between keywords in paper abstracts with cortical maps reported in the paper. We found that 27 out of 35 questions from our model had a reasonable match to a keyword available in Neurosynth (see Table S1), but we note that matches are often imprecise. We computed the correlations between the QA weights and the Neurosynth maps (mapped to subject coordinates, masked to the 10% best-predicted voxels by the QA model) and found reasonable agreement: 26 of the 27 correlations were positive and the mean correlation of 0.17 was statistically significant (𝑝 < 10^−5^, permutation test). (c) To validate the potentially new findings in the QA weights, we ran a follow-up experiment using generative causal testing (GCT). Specifically, we constructed synthetic stimuli designed to test each individual question and then measured the average response to these synthetic stimuli for subject S02. We computed the Pearson correlation between each GCT map and the QA weight and found that the mean correlation of 0.16 was significant (𝑝 < 10^−6^, permutation test). Masking the maps to include only well-predicted voxels improves the correlation with the GCT map; 8 out of 35 questions showed a statistically significant correlation for the top 1% of best-predicted voxels (𝑝 < 0.05, permutation test, FDR-corrected). (d) Finally, we assessed consistency across subjects by evaluating the correlation of the weights for each subject with the mean of the other two (averaged in MNI space then projected to the subject coordinates). The mean correlation of 0.30 was significant (𝑝 < 10^−6^, permutation test), and each individual correlation was also significant (𝑝 < 0.05, permutation test, FDR-corrected). The bottom row shows cortical selectivity map for visual comparison: (b) *place* projected onto S02, (c) GCT map for *location* in S02, and (d) QA weights for *location* in S03.

We first evaluated whether the QA-35 weights matched known selectivity maps from the literature. We compared the QA-35 weights with cortical maps from Neurosynth (*8*), a meta-analysis of the fMRI literature. Neurosynth used an automated pipeline that tagged voxel coordinates reported in neuroscience papers with keywords from the paper abstract, and then computed a selectivity map using an association test between each voxel and each keyword (see example in Fig. 3b). We manually searched through the keywords in Neurosynth and found a plausible match for 27 out of the 35 questions in QA-35 (see matches in Table S1). We note that matches were often imprecise—e.g. matches for *time* may relate to tasks related to *timing* rather than stimuli associated with the concept of *time*—biasing the correlation between the Neurosynth maps and QA-35 weights downwards. We then compared the Neurosynth maps with their corresponding QA-35 weights for each of three subjects by mapping the Neurosynth maps to subject coordinates, masking them to the 10% of best-predicted voxels by QA-35, and computing the correlation across voxels. The mean correlation of 0.17 across questions was significantly greater than zero (𝑝 < 10^−5^, permutation test), and 26 of the 27 individual correlations were positive, suggesting broad-scale agreement between the automated maps produced by QA-35 and the aggregated findings contained in Neurosynth.

We further assessed the QA-35 selectivity maps using targeted follow-up experiments. We conducted an fMRI experiment designed to measure selectivity for the underlying 35 questions using generative causal testing (GCT) (*9*). Specifically, we created two hours of synthetic natural stories, with each story paragraph designed to emphasize an individual question from QA-35. We then measured the average response to the paragraphs for each question in subject S02 and then computed the Pearson correlation between this average response and the QA-35 weights for that question (Fig. 3c). The whole-cortex correlation for 33 of the 35 questions was greater than zero and the mean correlation of 0.16 was significant (𝑝 < 10^−6^, permutation test). Well-predicted voxels showed better consistency with the GCT scores: the mean correlation when computed only over the best-predicted 1% of voxels (943 voxels) was 0.43, with 8 out of 35 questions yielding significant correlations (𝑝 < 0.05, permutation test with Benjamini-Hochberg false discovery rate correction (*20*)). To minimize the amount of new fMRI data to be collected, we reused fMRI responses to GCT synthetic stimuli for the subject S02 collected in a previous work (*9*), when there was a reasonable match between the QA question and the GCT synthetic stimuli (see Table S2).

Finally, we evaluated the consistency of the QA-35 weights across subjects (see example map for subject S03 in Fig. 3d). Specifically, for each subject we projected the weights for the other two subjects into MNI coordinates, averaged them, projected them to the original subject’s coordinates, and then measured the correlation with the original subject’s weights. The mean correlation of 0.30 was significantly greater than zero (𝑝 < 10^−6^, permutation test), as was each individual correlation (𝑝 < 0.05, permutation test, FDR-corrected). The questions with the highest inter-subject correlation often involved aspects of communication (e.g. *communication*, a *relationship*, a *social interaction*) or visual concepts (e.g. describe a *physical environment* or a *visual experience*).

These results demonstrate that the selectivity maps generated by QA-35 reflect known cortical function from the literature, are replicable in new stimuli, and are consistent across subjects.

## 5 QA encoding models for ECoG data

QA encoding models are highly effective at capturing fMRI responses to language, and here we investigate whether this success extends to another neuroimaging modality: electrocorticography (ECoG). We analyzed ECoG data from the Podcasts dataset (*21*), which contains responses from a total of 1,268 electrodes across nine subjects who each listened to the same 30 minute podcast. As was done with the fMRI data, we generated interpretable features for the stimuli using the questions in QA-35. As ECoG provides higher temporal resolution and sensitivity than fMRI, we generated the features at three contextual timescales: at the word level, with 1.5 seconds of context, and with 3 seconds of context. This yielded a total of 105 lexical and contextual QA features that we used to build our encoding model. We supplemented these QA features with low-level spectral features to capture both low-level acoustic and high-level semantic properties of the stimulus while remaining fully interpretable. From these concatenated features we built a model to predict ECoG responses from the high gamma frequency band at various time lags from word onset (*22*) (see Methods).

Fig. 4a shows the encoding performance of the resulting QA-35+Spectral model at the best performing lag for all 1,268 electrodes across all 9 subjects on a template brain. Many electrodes show strong performance with the QA features. The best prediction performance was found in left STG and near Broca’s area, with similar performance in the right hemisphere analogues of these regions.

**Figure 4:**
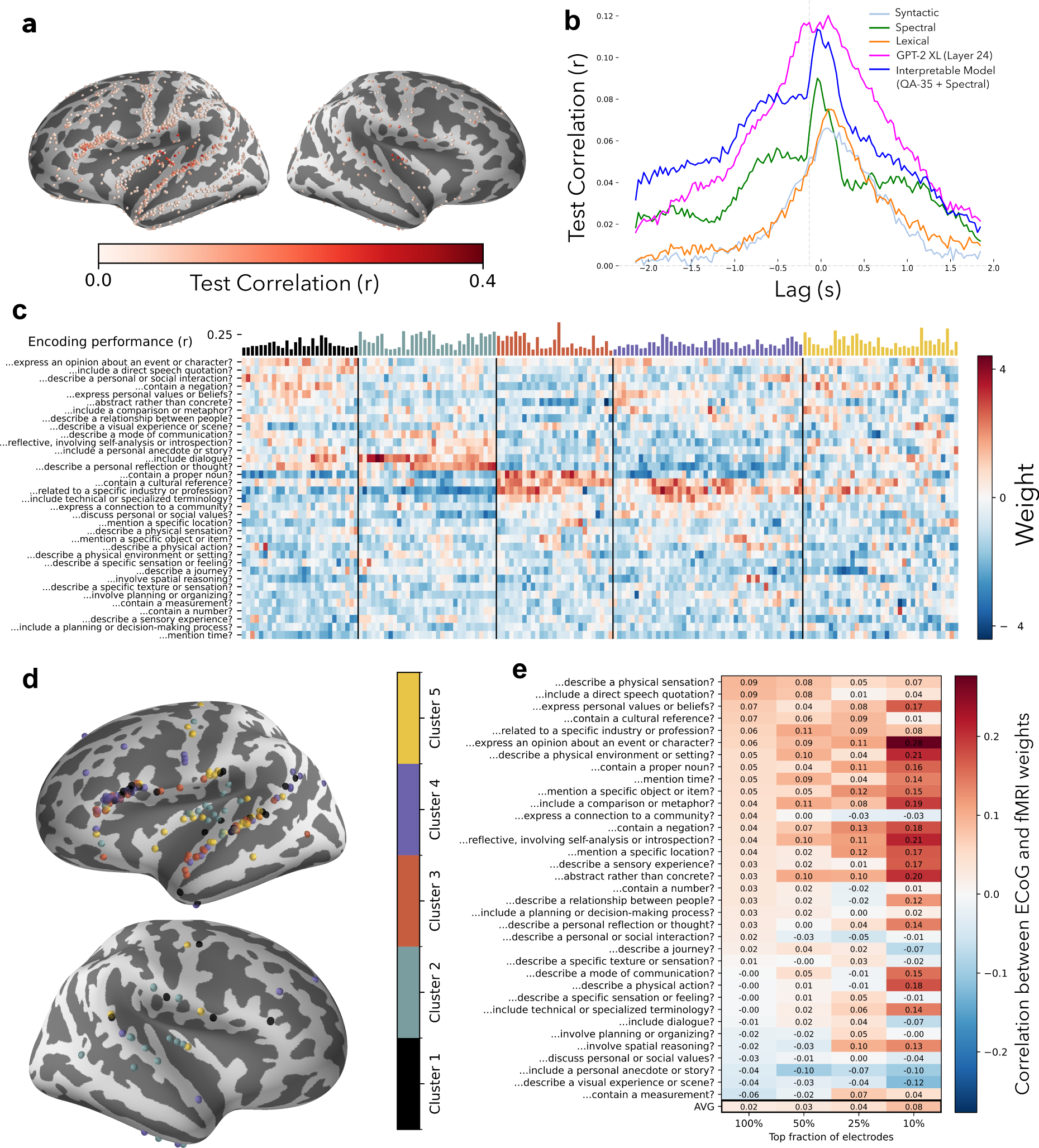
QA encoding models for ECoG data. To evaluate the applicability of QA encoding models to a different neuroimaging modality, we fit encoding models to ECoG responses in 1,268 total electrodes from 9 subjects listening to a 30-minute podcast. (a) For each electrode, we generated an interpretable encoding model by concatenating our 35-question QA encoding model from Fig. 1 with spectral features. We break down our model’s encoding performance, measured at the best performing overall lag, using use a joint template across all subjects. The best-predicted electrodes were found in left STG and near Broca’s area. (b) This model outperforms other interpretable encoding models and achieves similar performance to a non-interpretable baseline model from prior work, which uses GPT-2 XL hidden states as features (*pink*). (c) To understand selectivity trends across these electrodes, we select 166 well-predicted electrodes (see Methods) and visualize the encoding weights for each electrode. These electrodes form 5 high-level clusters with similar selectivity profiles. (d) We visualize these clusters and find that they display moderate spatial homogeneity, despite being aggregated across subjects. (e) We further investigate spatial selectivity by computing the spatial correlation of ECoG electrode weights with fMRI voxel weights for each question. We find statistically significant correlations when selecting different fractions of the best-predicted electrodes; a paired t-test between fMRI-ECoG correlations and permuted fMRI-ECoG correlations yields 𝑝 = (8.3 × 10^−4^, 3.6 × 10^−4^, 3.4 × 10^−5^, 2.7 × 10^−5^) for top electrode fractions (100%, 50%, 25%, 10%).

We compared the interpretable QA-35+Spectral model with various baseline encoding models from prior work on this dataset (*21*). We selected 166 electrodes based on their minimum prediction performance across all encoding models (see Methods) and performed the remaining analyses on the selected electrodes. Fig. 4b shows the performance of our interpretable model and baselines for each lag relative to word onset, averaged across the thresholded electrodes. As baselines, we built a model that only uses the spectral features, a syntactic model that includes simple part-of-speech / dependency parsing information, a black-box lexical embedding space (*en.core.web.lg*) and a black-box contextual embedding space from a large language model (*GPT-2 XL*, Layer 24) that was used in prior work (*21*). Our interpretable model achieved prediction performance that roughly matched the black-box LLM baseline at many lags and outperformed the other models, in spite of the fact that these questions were originally chosen for their ability to predict temporally coarser fMRI data. As all of the feature spaces contain information about the immediate temporal context, we were often able to predict neural response before word onset (*23, 24*).

To identify patterns of selectivity across electrodes, we visualized each electrode’s feature weights across the questions from QA-35 (Fig. 4c). Because the ECoG models included three separate features for each question that correspond to the three timescales used, we averaged weights across the timescales. Hierarchical clustering of these weights across electrodes illustrates various clusters of electrodes whose responses are driven by different types of questions. These clusters show some anatomical separation, as illustrated in Fig. 4d. For example, cluster 2, which strongly loads on the question *Does the input include dialogue?*, is largely located near primary auditory cortex. In contrast, cluster 3, which has high loadings on the *…contain a proper noun?* and *…contain a cultural reference?* questions, is spread throughout superior temporal gyrus and inferior frontal gyrus. This demonstrates the potential for these QA models, whose 35 features were not optimized for intracranial encoding, to reveal new patterns of response tuning.

Finally, to further test the consistency of our approach, we investigated the degree to which anatomical patterns across the two modalities aligned. Fig. 4e shows the cross-modality correlations between the fMRI weights and the ECoG weights for each question. All fMRI and ECoG weights were first mapped to the same FreeSurfer fsaverage template coordinates (*25*). fMRI weights were mapped from volumetric coordinates to vertices on the template, and then a distance-weighted average of the nearest 10 vertices was used as a proxy for the fMRI weight for each electrode location. The correlation between these interpolated weights from fMRI and ECoG was then directly computed. Overall, the correlation between fMRI and ECoG maps is significantly greater than zero (𝑝 = 8.3 × 10^−4^; paired t-test between the correlations of the fMRI and ECoG maps with and without permutation), with stronger correlation when only the best-performing electrodes are considered. The few negatively correlated features, such as *…describe a journey?* or *…describe a visual experience or scene?*, likely reflect undersampling of the relevant cortical areas in the ECoG dataset. The significant correlations between the spatial patterns uncovered by fMRI and ECoG QA encoding models demonstrate that the uncovered patterns are robust across datasets and modality.

## 6 Discussion

Many prior studies have demonstrated the predictive power of black-box encoding models in language neuroscience across modalities such as fMRI, ECoG, EEG, and MEG (*1,4,5,6,7*). However, these models have been criticized for their inability to produce accurate and reliable explanations (*26, 27*) and attempts to explicitly interpret these models have been limited (*28, 29, 30*). Our work addresses this criticism by directly constructing a fully interpretable encoding model using LLM-generated annotations.

The methods here are related to work that has used LLMs as tools for data annotation (*31*) in domains such as medicine (*32, 33*), finance (*34*), and general natural-language processing (*35, 36*). They also share a looser connection with interpretable text representations, which are often built by averaging word-level embeddings (*15, 37, 38*) or ngram-level embeddings (*39, 40*). In contrast, we use LLMs as tools not just for annotating and representing data, but for formulating and testing qualitative theories.

This distinction provides an important scientific benefit: it enables empirical evaluation of theory’s predictive power and iteration on theories to refine their accuracy and detail. The process of generating these theories may be aided by methods for the post-hoc interpretation of black-box models (*41,42,11*) that can generate refined descriptions of encoding model behavior in specific regions or domains of interest. QA encoding models can efficiently evaluate the predictive power of these theories and guide the design of follow-up experiments that causally test these theories, e.g. through generative causal testing (*9*).

While effective, QA encoding models have two major limitations. First, QA encoding models are computationally intensive, requiring an LLM call for every question at every timepoint. This is often prohibitively expensive, but will likely become more feasible as LLM inference costs continue to decrease. Additionally, a single LLM model can be distilled to answer multiple questions simultaneously when the questions are predefined (*36, 12*). Second, QA encoding models rely on the underlying LLM’s ability to faithfully answer the given yes-no questions (see evaluation in the Methods). If an LLM is unable to accurately answer the questions, it compromises the interpretability of the QA encoding models. Thus, QA encoding models require the use of strong LLMs that can accurately answer the set of chosen questions. This challenge will also likely diminish as LLM capabilities improve.

The work here opens many avenues for future research. One approach can explore using QA encoding models for various input modalities. Modern multimodal foundation models can answer textual questions about diverse modalities, including audio and images (*43*), enabling annotation for a broader range of stimuli. Additionally, QA encoding models can be readily applied to emerging scanning modalities (e.g. 7T laminar fMRI or NeuroPixels), which may reveal selectivity patterns that were previously undetectable. Finally, while we take an initial step towards characterizing cortex-wide selectivity map, QA encoding models can easily be tailored to answer more targeted theories about selectivity in the brain. In the appendix we include example vignettes illustrating this application–e.g. using QA encoding models to identify monosemantic voxels (Section S2.1), describe distinctive characteristics of the language network (Section S2.2), and decoding the answers to different questions (Section S2.3).

QA encoding models offer a general method for evaluating theories through data-driven predictions. In this paper, we demonstrate how modern LLMs can enhance our understanding of cognitive brain processes. Through simple prompt-based methods, we annotate large natural datasets to recover selectivity theories that traditionally required decades of painstaking controlled experiments and pave the way for many similar future discoveries. More broadly, the core concept of QA encoding models–using foundation models to operationalize qualitative theories about the natural world–has applications well beyond neuroscience. QA encoding models will help advance scientific discourse toward increasingly nuanced, data-supported explanations.

## Author contributions

C.S., R.J.A., and A.G.H. led the experimental conception and design, with input from all authors. C.S. led the fMRI data analysis and R.J.A. led the ECoG data analysis. S.G. and R.J.A. collected fMRI data. S.G. led the comparisons with Neurosynth and G.M. helped with ECoG evaluation and analysis. Writing was primarily handled by C.S., R.J.A., and A.G.H., with editing from all authors. N.M, J.G., and A.G.H. provided funding and supervision for the project.

## Data and materials availability

All newly collected fMRI data will be made publicly available upon acceptance. Data for fitting encoding models and generating explanations is available on OpenNeuro: openneuro.org/datasets/ds003020 (fMRI) and openneuro.org/datasets/ds005574/versions/1.0.2 (ECoG).

Code for running all experiments (as well as applying QA encoding models in new settings) is available on Github at github.com/microsoft/automated-brain-explanations. Code uses python 3.10, huggingface transformers 4.29.088–100, sklearn 1.2.040, and OpenAI API GPT-4 (*gpt-4-0125-preview*).

## Supplementary materials

### S1 Materials and methods

**fMRI data collection** This study uses two MRI datasets: one for fitting encoding models and a second for evaluating cortical selectivity maps. The original set is described and made publicly available in previous work (*13, 44*), while details of the newly collected second dataset are described here. Functional magnetic resonance imaging (fMRI) data were collected from a single subject (S02) in the original set as stories were visually presented at approximately conversational cadence. We collected two hours of data corresponding to 9 stories. Participants passively listened to the stories without making any responses. All subjects were healthy and had normal hearing. The experimental protocol was approved by the Institutional Review Board at the University of Texas at Austin and written informed consent was obtained.

All MRI data were collected on a 3T Siemens Skyra scanner at the University of Texas at Austin using a 64-channel Siemens volume coil. Functional scans were collected using a gradient echo EPI sequence with repetition time (TR) = 2.00 s, echo time (TE) = 30.8 ms, flip angle = 71°, multi-band factor (simultaneous multi-slice) = 2, voxel size = 2.6mm x 2.6mm x 2.6mm (slice thickness = 2.6mm), matrix size = 84×84, and field of view = 220 mm. Anatomical data were collected using a T1-weighted multi-echo MP-RAGE sequence with voxel size = 1mm x 1mm x 1mm following the Freesurfer morphometry protocol (*25*).

**fMRI data preprocessing** All functional data were motion corrected using the FMRIB Linear Image Registration Tool (FLIRT) from FSL 5.0. FLIRT was used to align all data to a template that was made from the average across the first functional run in the first story session for each subject. These automatic alignments were manually checked for accuracy.

Low frequency voxel response drift was identified using a 2nd order Savitzky-Golay filter with a 120 second window and then subtracted from the signal. To avoid onset artifacts and poor detrending performance near each end of the scan, responses were trimmed by removing 20 seconds (10 volumes) at the beginning and end of each scan. This process eliminated the 10-second silent period and the first and last 10 seconds of each story. The mean response for each voxel was subtracted and the remaining response was scaled to have unit variance.

**Generating questions for QA encoding models** To generate the questions underlying QA encoding models, we prompted GPT-4 (*45*) (*gpt-4-0125-preview*) with 6 prompts that aimed to elicit semantic information that was useful for predicting fMRI responses (precise prompts in Section S3.1). This included directly asking the LLM to use its knowledge of neuroscience, to brainstorm semantic properties of narrative sentences, to summarize examples from the input data, and to generate questions similar to single-voxel explanations found in a prior work (*11*). Many of the prompts included examples of diverse, reasonable questions, incorporating the author’s domain knowledge. After deduplication, this process yielded 606 questions (see all the questions on Github).

**Extracting QA features** For answering questions, we took the mean of the answers from Mistral-7B (*46*) (*mistralai/Mistral-7B-Instruct-v0.2*), LLaMA-3 8B (*47*) (*meta-llama/Meta-Llama-3-8B-Instruct*) with two prompts, and GPT-4 (*gpt-4-0125-preview*). QA features were extracted using 64 AMD MI210 GPUs, each with 64 gigabytes of memory. See all prompts in Section S3.1.

If an LLM is unable to accurately answer the questions, this compromises the interpretability of the QA encoding models. Thus, QA encoding models require the use of high-performing LLMs, and the set of chosen questions must be accurately answered by these models. Section S3.4 provides an analysis of the question-answering accuracy of different LLMs and finds that the LLMs used in this study can reliably answer the queried questions.

**Regression modeling** Each subject’s fMRI data consists of approximately 100,000 voxels; we pre-processed it by running principal component analysis (PCA) and extracting the coefficients of the top 100 components for each TR. We then fit ridge regression models to predict these 100 coefficients. We still evaluated the models in the original voxel space (by applying the inverse PCA mapping and measuring the correlation between the response and prediction for each voxel in the test set). To handle temporal sampling, we followed the approach in prior works (*15, 1*); an embedding was computed at the timepoint where each word occurred in the input story, and these embeddings were interpolated using Lanczos resampling. Embeddings at each timepoint were computed from the ngram consisting of the 10 preceding words. We selected the best-performing hyperparameters via cross-validation on 5 time-stratified bootstrap samples of the training set. The best ridge parameters were chosen from 12 logarithmically spaced values between 10 and 10,000. To model temporal delays in the fMRI signal, we selected between adding 4, 8, or 12 time-lagged duplicates of the stimulus features. After fitting and evaluating the encoding model, we averaged across the temporal delay dimension of the weights for visualization and comparison (e.g. Fig. 3).

**Selecting stable QA features** We selected a compact feature set using stability selection (*14*). Specifically, we fit multi-task Elastic net with 10 logarithmically spaced regularization parameters ranging from 10^−3^ to 1 using the *MultiTaskElasticNet* class from scikit-learn (*48*). We randomly sampled the training dataset by 50% five times and fit the Elastic net model to each of the sets. We then selected only the only the stable features for each regularization parameter, i.e. those that were consistently chosen across all five training sets for a given regularization parameter. After feature selection, we refit the Ridge regression model using only the selected features.

**Baselines** We compared QA encoding models to Eng1000, an interpretable baseline developed in the neuroscience literature specifically for the task of predicting fMRI responses from narrative stories (*15*). Each element in an Eng1000 embedding corresponds to a co-occurence statistic with a different word, allowing full interpretation of the underlying representation in terms of related words. We additionally compared QA encoding models to embeddings from BERT (*49*) (*bert-base-uncased*) and LLaMA models (*17,47*). For each subject, we swept over 5 layers from LLaMA-2 7B (*meta-llama/Llama-2-7b-hf*, layers 6, 12, 18, 24, 30), LLaMA-2 70B (*meta-llama/Llama-2-70b-hf*, layers 12, 24, 36, 48, 60), and LLaMA-3 8B (*meta-llama/Meta-Llama-3-8B*, layers 6, 12, 18, 24, 30). We selected the best layer using cross-validation and then reported its test performance.

**Significance testing of cortical map correlations** In Section 4, we performed permutation tests to compare the correlation between different cortical maps. All permutation tests were performed relative to a null distribution of 2,000 randomly selected response TRs from the training data for each subject. Since these responses were recorded during a passive listening task, they generally represent semantic processing information present in the stimulus while preserving correlation structure among the voxels.

**Expert survey** To evaluate the findings in the QA selectivity maps against expert opinion, we conducted an anonymous survey asking researchers to provide their judgment of each question’s importance for predicting brain responses to language using a five-point Likert scale (e.g. 1 = “Not at all important”, 3 = “Moderately important”, 5 = “Extremely important”). We included the 35 questions in QA-35. From the remain 571 questions, we additionally included 5 of the 15 most highly predictive questions and 5 of the 15 most poorly predictive questions, manually selected to be diverse (see selected questions in Fig. S1(c)). The exact survey prompt was “Below, we have listed 45 potential properties of language stimuli. For each property, we would like you to rate how important that property is for predicting brain responses to language. Specifically, for each property, please rate the degree to which knowing that a stimuli possesses that property would be useful for you to determine which brain regions might be responsive to that stimuli.”

The survey was sent out to four relevant mailing lists (cvnet, visionscience, and computational neuroscience) and yielded 12 responses. The survey protocol was approved by the Institutional Review Board at Microsoft Research and written informed consent was obtained.

**ECoG data details** The Podcast ECoG dataset (*21*) comprises electrocorticography (ECoG) recordings from nine epilepsy patients who listened to a 30-minute audio podcast. We summarize the dataset and its preprocessing details here (further details can be found in the dataset paper). The 1,330 electrodes in the initial dataset are filtered down to 1,268 electrodes by removing electrodes with poor localization or noisy power spectrum density. All electrode visualizations were plotted on the *fsaverage* template. We utilized the high-gamma band power of the signal for all of our analyses, extracted by applying a Butterworth band-pass infinite impulse response filter from 70Hz to 200Hz.

**ECoG modeling details** We predicted ECoG responses from the high gamma frequency band at various time offsets from word onset following a prior work (*22*). Specifically, for each of 128 equally-spaced lags in between −2 and 2 second offsets from word onset, we predicted the electrode response as a function of the QA features. Ridge regression was used to produce a linear model from the stimulus features to the responses for each (electrode, lag) pair. Since the amount of training data is small, model performance was estimated using 5-fold cross-validation of the dataset, training on 80% of the data and testing on the remaining 20% for each of 5 folds. Within each fold, the regularization parameter 𝛼 of this regression was estimated using boostrapped cross-validation.

For follow-up analysis, we selected electrodes only if they had a best-lag encoding performance of at least 𝑟 = 0.06 across all examined feature spaces (*syntactic*, *spectral*, *En.core.web.lg*, *GPT-2 XL*, *Interpretable*). This performance threshold was selected using the elbow method (see Fig. S10). The selection process yielded 166 electrodes.

### S2 Additional analyses using QA encoding models

#### S2.1 Evaluating single-question QA encoding models

**Figure S1:**
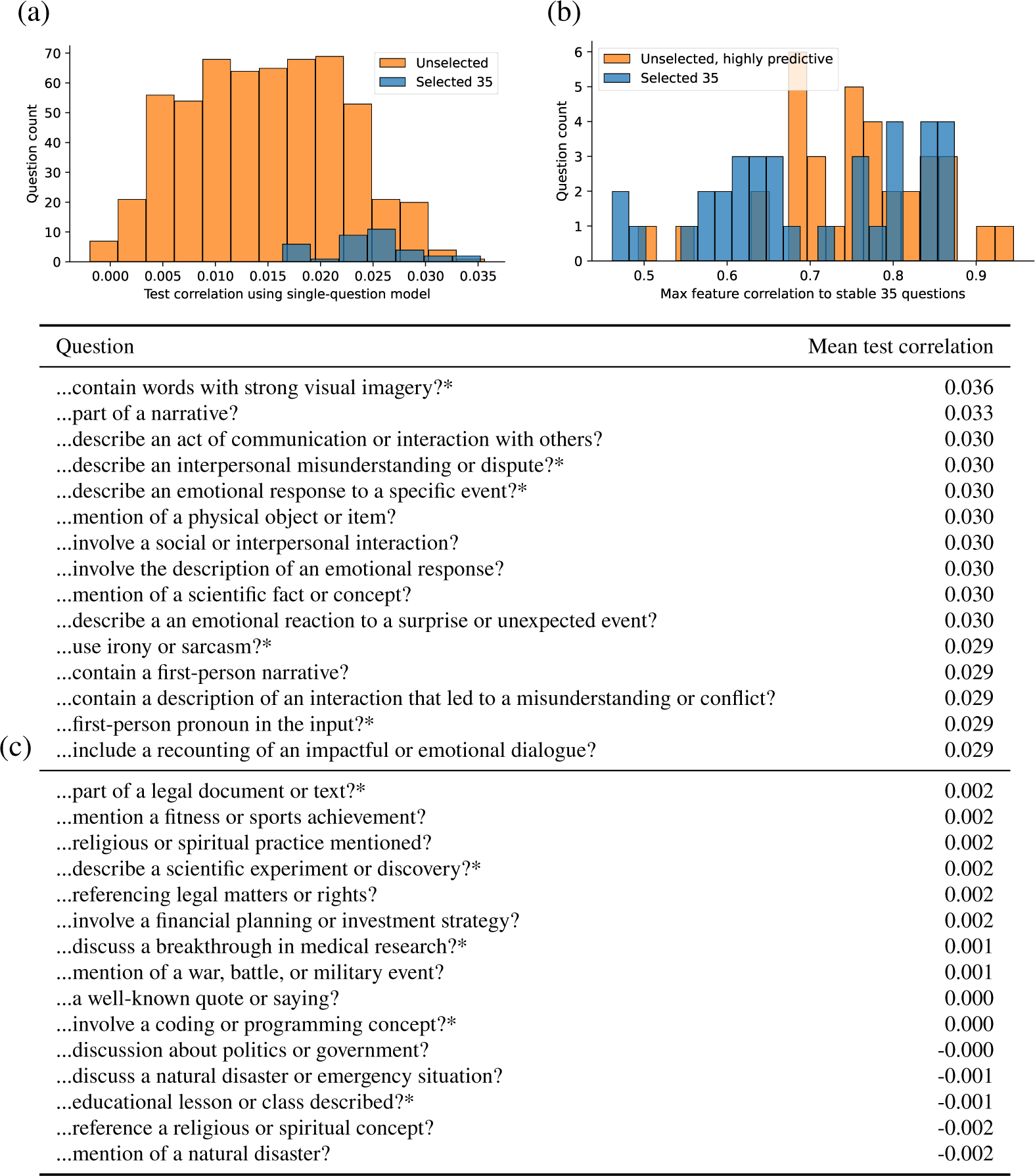
Single-question QA encoding models. (a) We computed the prediction performance (mean test correlation) when building QA encoding models using the features extracted by only a single question. The single-question models corresponding to the 35 selected questions yielded stronger performance (mean correlation of 0.025) than the remaining 571 questions (mean correlation of 0.015). (b) The unselected but highly predictive questions often were highly related to one of the 35 selected questions. To demonstrate this, we computed the maximum correlation between the features extracted for a question and the features extracted for each of the 35 selected questions (excluding the question itself). For the 35 unselected questions with the highest individual predictive performance (orange), the mean maximum correlation was 0.70. This is higher than the mean maximum correlation for the 35 selected questions (blue), which was 0.51. (c) We show the 15 most predictive single questions that were not selected in QA-35 as well as the 15 least predictive single questions. Questions marked with an asterisk were used as part of the external expert survey described in the Methods.

**Figure S2:**
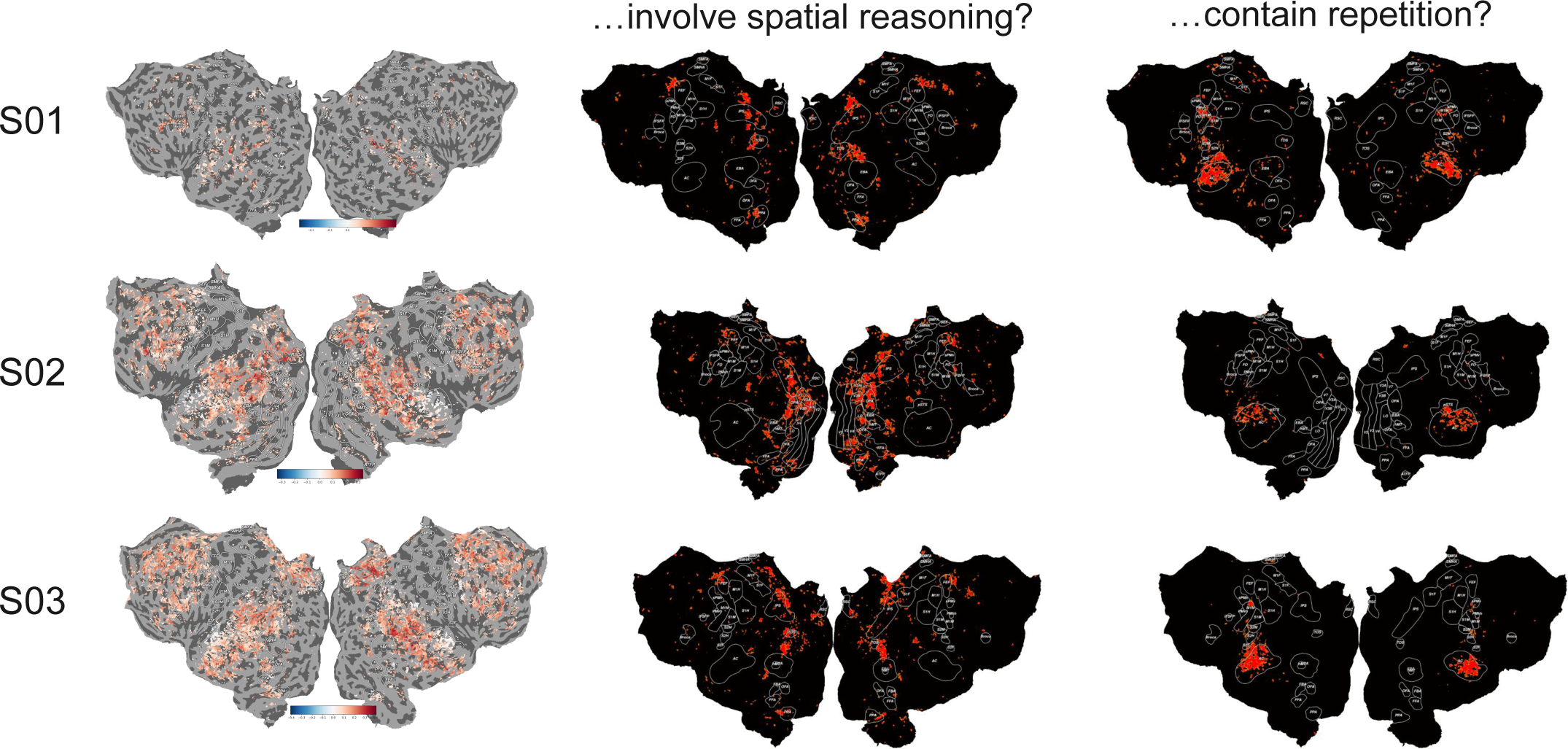
Monosemanticity in fMRI voxels. We fit monosemantic encoding models to each voxel by selecting the single-question encoding model that achieved the best cross-validation performance on the training set. The left column displays the difference between the test correlation for the full 35-question model and a monosemantic model. We find that individual questions can predict a small fraction of voxels reasonably well, but most voxels are polysemantic–i.e. incorporating more QA features improves performance. The well-predicted monosemantic regions (white) often correspond to one of two questions: either *involving spatial reasoning* (middle column) or *containing repetition* (right column).

#### S2.2 Characterizing language network voxels with QA encoding models

**Figure S3:**
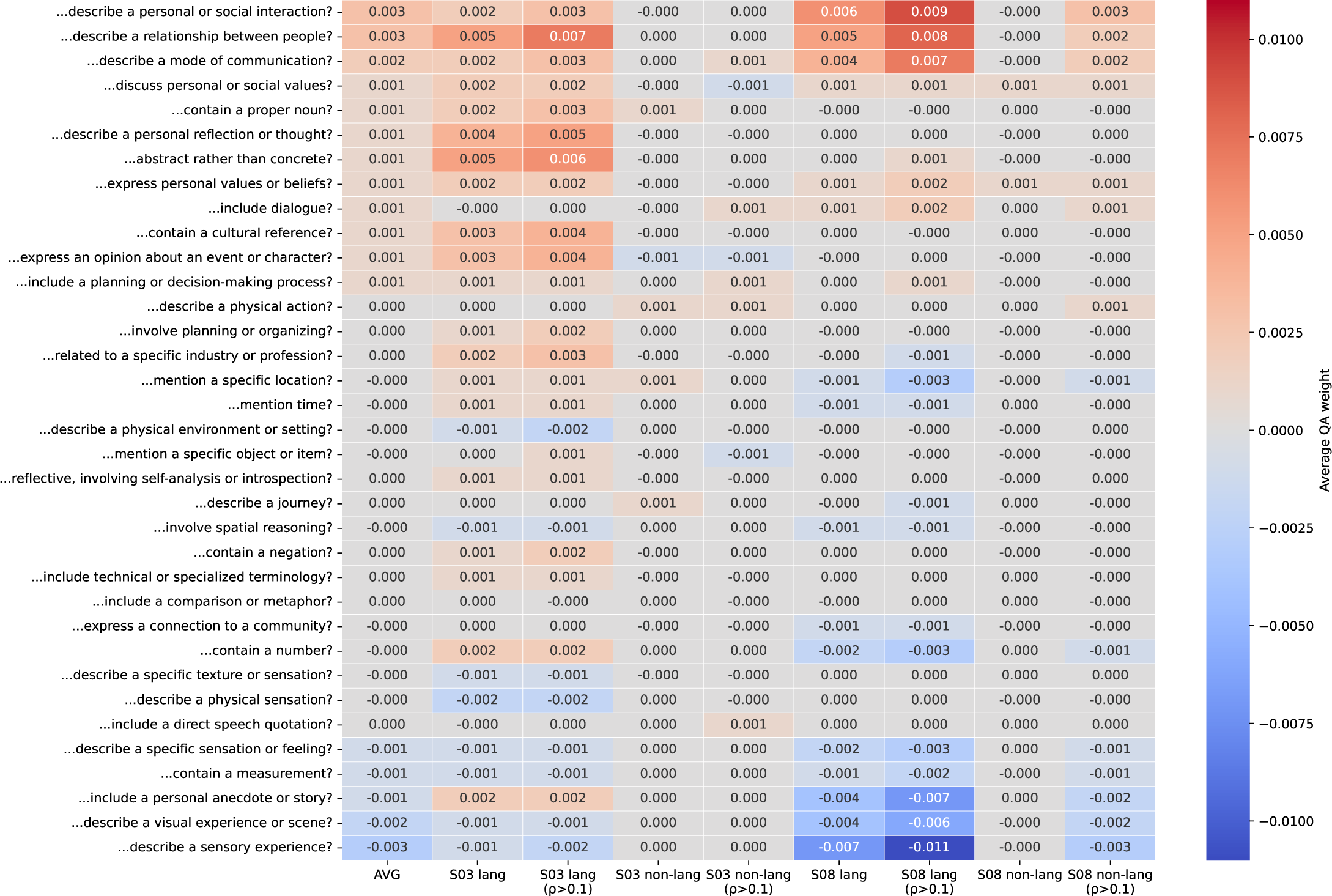
Characterizing voxels in the language network. Language network voxels–i.e., voxels that respond significantly more to sentences than non-words–were identified using an auditory language localizer (*50*) task. These voxels were identified as any cortical voxel that passed a one-sided t-test with a significance threshold of 𝑝 <= 0.001 (uncorrected). We then evaluated the overall selectivity of the language network by computing the average QA weight for language-network voxels compared to non-language-network voxels. We further filtered voxels based on whether they were well-predicted by the QA-35, meaning they achieved a test correlation greater than 0.1. Across two subjects, a handful of questions showed distinct differences between language-network and non-language-network voxels.

#### S2.3 Decoding QA features

While the main results focused so far on encoding models, QA features can instead be used as targets for semantic decoding models, i.e. models that predict the QA answer at a particular timepoint from the recorded brain responses. These decoding models may be useful in different scenarios than encoding models, such as improving brain-machine interfaces (*51, 52*).

To build decoding models in the fMRI data, we first constructed binary QA labels for each TR by temporally downsampling the binary answers for each 10-gram and thresholding the result at a z-score of 1. We then predicted this label independently for each question using logistic regression. As inputs to the logistic regression, we used the voxel responses recorded in the four TRs following the decoded TR. To make comparisons clearer, we subsampled the data for each question to equally balance the labels in both the training and testing set.

**Figure S4:**
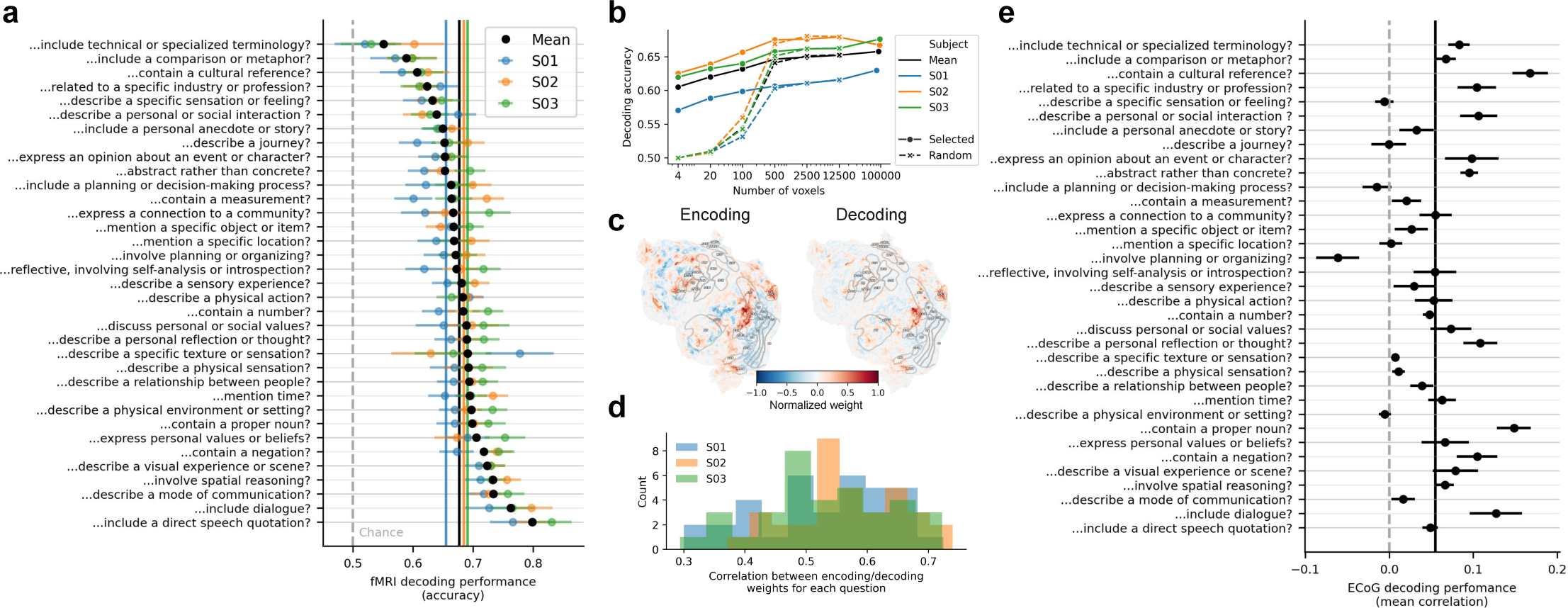
Decoding individual QA features from fMRI. (a) We linearly decoded the answers to 35 stable questions for each TR from the four subsequent TRs. Decoding accuracy for all questions was above chance (chance was set at 0.5 by undersampling each question to have 50% yes answers). The three best-decoded questions involve speech and communication. (b) We then decoded the answers using only the responses from voxels that had the largest positive initial decoding weights. This masking strategy yielded fairly high decoding accuracy compared to decoding from randomly selected voxels, and even achieved reasonably high decoding accuracy using only 4 voxels. (c) We then compared the weights for encoding and decoding. Example weights for a question about *locations* are shown for subject S02’s left hemisphere, both again emphasizing well-known location-selective regions such as RSC, PPA, and OPA. (d) The weights for encoding and decoding are generally quite similar. The similarity (averaged over the three subjects) is statistically significant for each question (𝑝 < 0.05, permutation test, FDR-corrected). (e) We performed an analogous decoding analysis using ECoG data and decoded the responses to each question at a much finer timescale. Most questions were decoded above chance, measured using the mean correlation of the decoded label and averaged over 9 subjects. All error bars show the standard error of the mean.

Fig. S4a shows that all questions were decoded above chance for the three subjects, with a mean decoding accuracy of 0.677. The best-decoded questions often involved communication, e.g. a *speech quotation*, *dialogue*, or a *mode of communication*. To test whether decoding could be performed from a restricted set of voxels, we repeated the decoding experiment using only the responses from voxels that had the largest positive decoding weights. We found that this masking strategy achieved fairly high decoding accuracy compared to decoding from randomly selected voxels (Fig. S4b), and even achieved reasonably high decoding accuracy using as little as 4 voxels (mean decoding accuracy 60.5%).

We then compared the QA weights for encoding and decoding the same question. Interpreting weights from a decoding model can be difficult: even if a concept is reflected in a voxel, it may not be uniquely reflected in the voxel and therefore assigned a low decoding weight (*53, 54*). Nevertheless, we observed strong alignment between the encoding and decoding weights for each question (see example selectivity for the question *Does the input mention a specific location?* in Fig. S4c). The mean Pearson correlation between encoding and decoding weights for each question was 0.545 (significant with 𝑝 < 10^−6^, permutation test) and the average correlation across 3 subjects was significant for every question (𝑝 < 0.05, permutation test, FDR-corrected). We found that large decoding weights were often concentrated in auditory cortex, more so than encoding weights (see averages in Fig. S5).

We similarly decoded the labels for each question in the ECoG dataset at each timepoint (see Methods). We were again able to decode most questions above chance (Fig. S4e). Taken together, these decoding results further demonstrate that the sparse 35 questions accurately capture major dimensions of semantic and cognitive selectivity on cortex.

**Figure S5:**
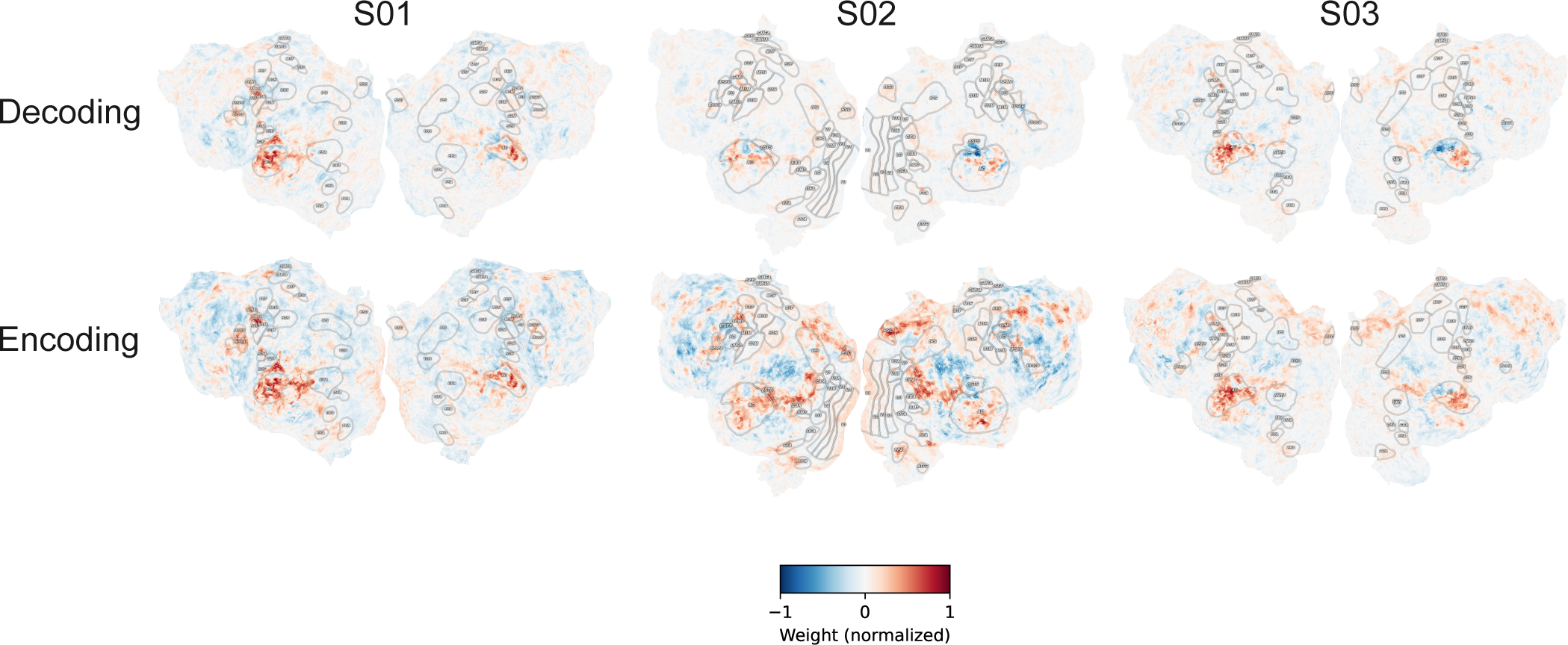
Average decoding and encoding weights across questions. Auditory cortex tends to have the largest values, especially for decoding. This is likely because auditory cortex captures information regarding the local word rate and the QA features are often zero when there are no (or few) words spoken in a TR, a consistent trend across all questions.

#### S2.4 Extended survey results

**Figure S6:**
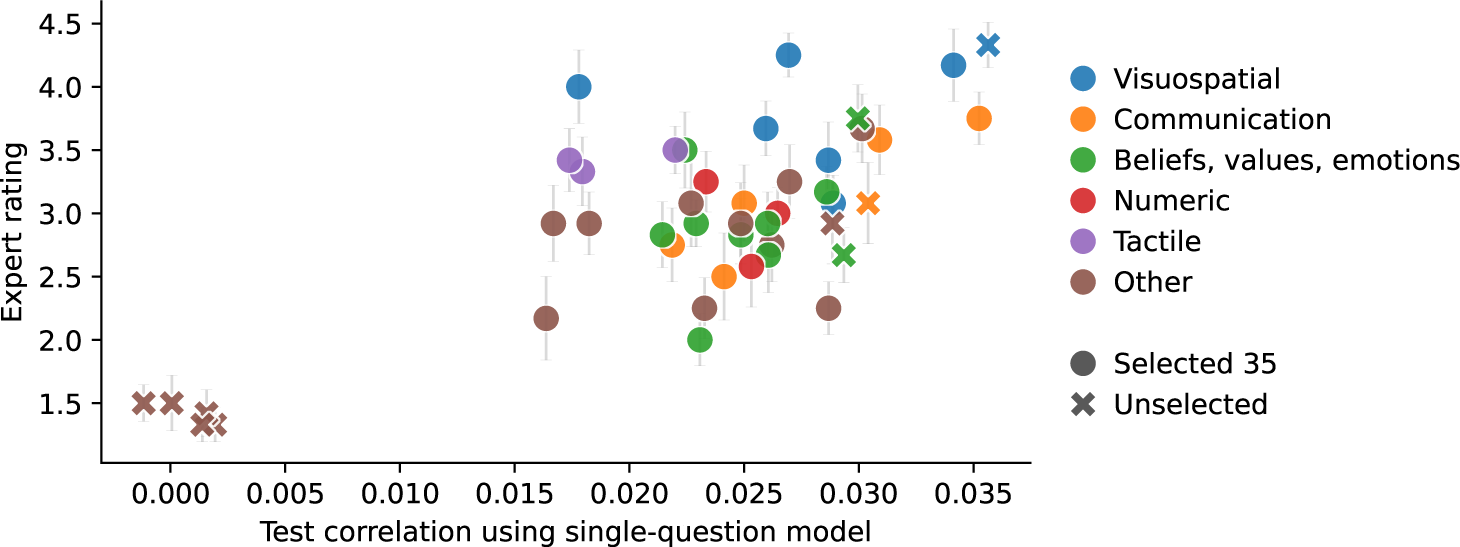
To evaluate the findings of the QA-35 selectivity maps against expert opinion, we conducted an anonymous survey asking researchers to rate questions based on how important they are for predicting brain responses to language using a five-point Likert scale (1 = “Not at all important”, 5 = “Extremely important”). We included the 35 questions in QA-35 along with 5 highly predictive and 5 poorly predictive questions from the remaining 571 questions. The survey was sent out to four relevant mailing lists and yielded 12 responses; see full survey details in the Methods. The experts clearly identified the 5 poorly predictive questions (mean rating 1.14) and were moderately successful at ranking the remaining questions (mean correlation 0.37, even after excluding the 5 worst-predicting questions). The raters show mild but significant inter-rater agreement with a Fleiss’ kappa of 0.098 (𝑝 < 10^−3^, permutation test).

### S3 Extended data and methods details

#### S3.1 Prompting details

See the general prompts used for eliciting questions below. Note that *{{examples}}* was filled in with 5-10 examples representing diverse concepts, e.g. *Does the input contain a number?*, *Does the input mention laughter?*, *Is hair or clothing mentioned in the input?*. See the exact examples in the Github repo.

**Question generation prompt 1** *Generate a bulleted list of 500 diverse, non-overlapping questions that can be used to classify an input based on its semantic properties. Phrase the questions in diverse ways.*

*Here are some example questions:*
*{{examples}}*
*Return only a bulleted list of questions and nothing else*

**Question generation prompt 2** *Generate a bulleted list of 100 diverse, non-overlapping questions that can be used to classify sentences from a first-person story. Phrase the questions in diverse ways.*

*Here are some example questions:*
*{{examples}}*
*Return only a bulleted list of questions and nothing else*

**Question generation prompt 3** *Generate a bulleted list of 200 diverse, non-overlapping questions that can be used to classify sentences from a first-person story. Phrase the questions in diverse ways.*

*Here are some example questions:*
*{{examples}}*
*Return only a bulleted list of questions and nothing else*

**Question generation prompt 4** *Based on what you know from the neuroscience and psychology literature, generate a bulleted list of 100 diverse, non-overlapping yes/no questions that ask about properties of a sentence that might be important for predicting brain activity.*

*Return only a bulleted list of questions and nothing else*

**Question generation prompt 5** *# Example narrative sentences {{example sentences from dataset}}*

*# Example yes/no questions*
*{{example questions already asked}}*

Generate a bulleted list of 100 specific, non-overlapping yes/no questions that ask about aspects of the example narrative sentences that are important for classifying them. Focus on the given narrative sentences and form questions that combine shared properties from multiple sentences above. Do not repeat information in the example questions that are already given above. Instead, generate complementary questions that are not covered by the example questions. Return only a bulleted list of questions and nothing else.

**Question generation prompt 6** *Generate more diverse questions that may occur for a single sentence in a first-person narrative story*

**Question answering standard prompt**

~~~
<User>: Input text: \{example\}\n
Question: \{question\}\n
Answer with yes or no, then give an explanation.}
~~~

**Question answering few-shot prompt**

~~~
<System>: You are a concise, helpful assistant.\n
<User>: Input text: and i just kept on laughing because it was so
Question: Does the input mention laughter?
Answer with Yes or No.
<Assistant>: Yes
<User> Input text: what a crazy day things just kept on happening
Question: Is the sentence related to food preparation?
Answer with Yes or No.
<Assistant>: No
<User> Input text: i felt like a fly on the wall just waiting for
Question: Does the text use a metaphor or figurative language?
Answer with Yes or No.
<Assistant>: Yes
<User> Input text: he takes too long in there getting the pans from
Question: Is there a reference to sports?
Answer with Yes or No.
<Assistant>: No
<User> Input text: was silent and lovely and there was no sound except
Question: Is the sentence expressing confusion or uncertainty?
Answer with Yes or No.
<Assistant>: No
<User> Input text: \{example\}
Question: \{question\}Answer with Yes or No.
<Assistant>:
~~~

#### S3.2 Cortex map matching details

**Table S1:** Matches between the questions in QA-35 and Neurosynth keywords.

| Question | Neurosynth keyword |
| --- | --- |
| ...contain a measurement? | arithmetic |
| ...contain a number? | arithmetic |
| ...describe a specific texture or sensation? | sensation |
| ...involve planning or organizing? | planning |
| ...contain a negation? | negative |
| ...contain a proper noun? | nouns |
| ...describe a personal or social interaction ? | social-interaction |
| ...describe a personal reflection or thought? | thoughts |
| ...describe a physical action? | action |
| ...describe a physical sensation? | touch |
| ...describe a relationship between people? | social |
| ...describe a specific sensation or feeling? | sensation |
| ...describe a visual experience or scene? | visual-information |
| ...express an opinion about an event or character? | judgments |
| ...include a direct speech quotation? | communication |
| ...include a personal anecdote or story? | personal |
| ...include dialogue? | communication |
| ...describe a physical environment or setting? | location |
| ...discuss personal or social values? | social |
| ...involve spatial reasoning? | spatial |
| ...mention a specific object or item? | object |
| ...mention a specific location? | place |
| ...describe a mode of communication? | communication |
| ...include a planning or decision-making process? | planning |
| ...abstract rather than concrete? | abstract |
| ...reflective, involving self-analysis or introspection? | thoughts |
| ...mention time? | time-task |
| ...related to a specific industry or profession? | NO MATCH |
| ...express personal values or beliefs? | NO MATCH |
| ...describe a sensory experience? | NO MATCH |
| ...include technical or specialized terminology? | NO MATCH |
| ...contain a cultural reference? | NO MATCH |
| ...include a comparison or metaphor? | NO MATCH |
| ...express a connection to a community? | NO MATCH |
| ...describe a journey? | NO MATCH |

**Table S2:** Imprecise matches for generative causal testing experiments. We used generative causal testing to evaluate the questions underlying the fitted QA encoding models. While we built precise matches for 31 of the 35 questions, 4 questions have imprecise matches as they were matched with stimuli from the original GCT study (*9*) rather than constructed specifically for the study here.

| Question | GCT driving explanation (imprecise) |
| --- | --- |
| Does the sentence involve the mention of a specific object or item? | Descriptive elements of scenes or objects |
| Does the input include a comparison or metaphor? | abstract descriptions |
| Does the sentence express a sense of belonging or connection to a place or community? | Relationships |
| Does the text describe a journey | Spatial positioning and directions |

#### S3.2 Selectivity maps

**Figure S7:**
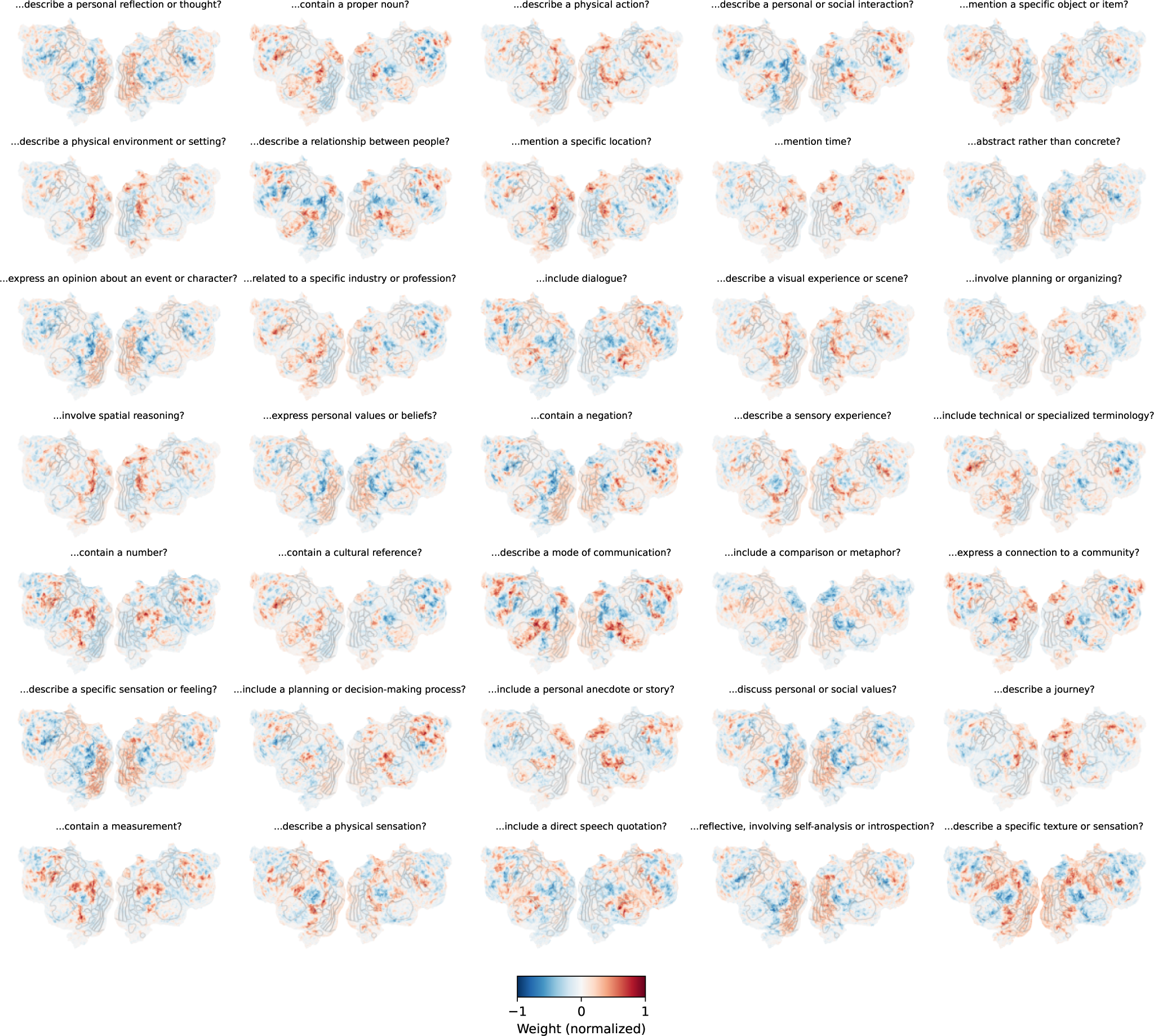
QA encoding weights for each question in a single subject (S02).

**Figure S8:**
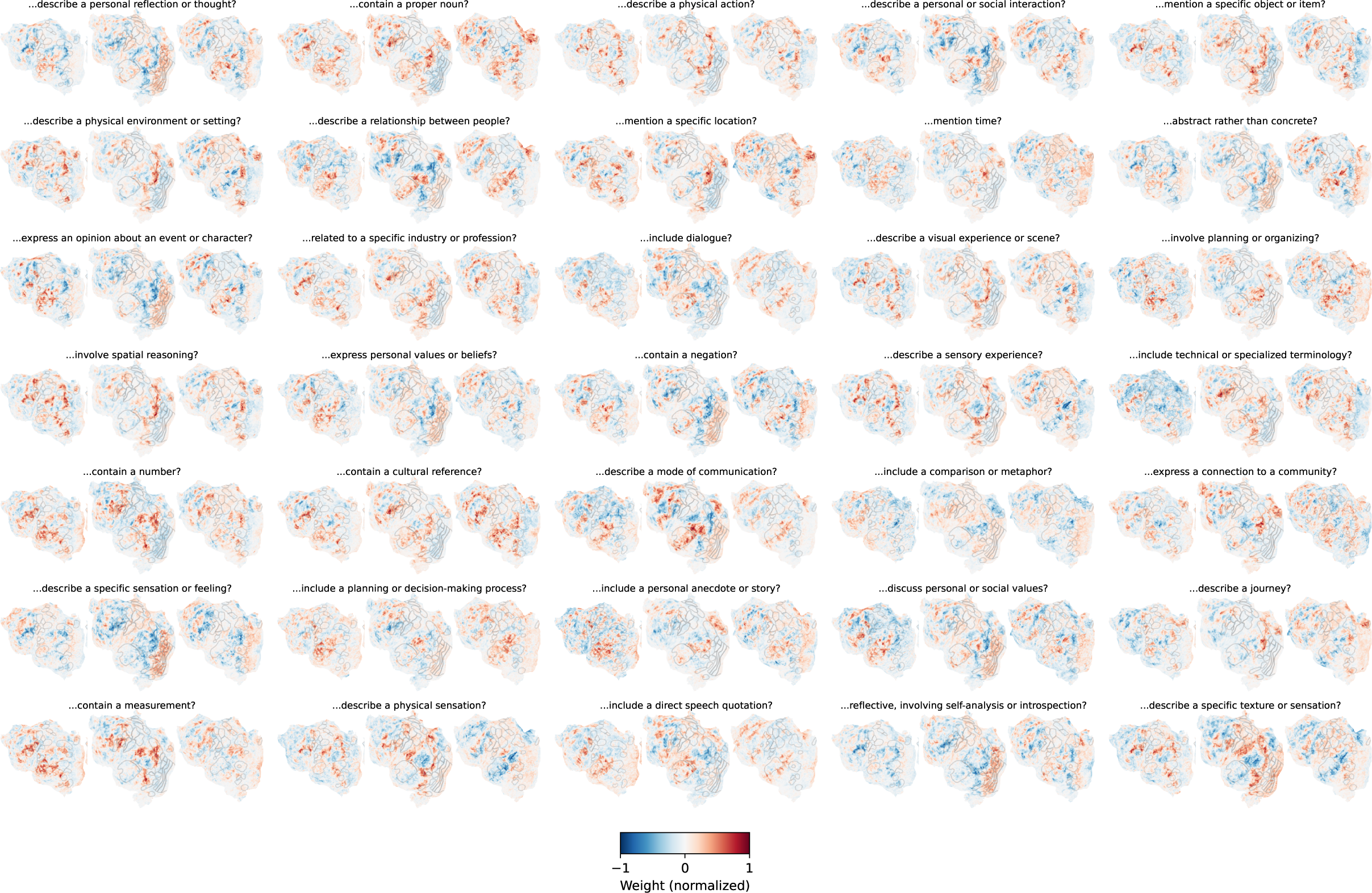
QA encoding weights for each question in QA-35 in left hemisphere for three subjects (S01, S02, and S03 from left to right).

#### S3.4 Evaluating question-answering faithfulness

**Figure S9:**
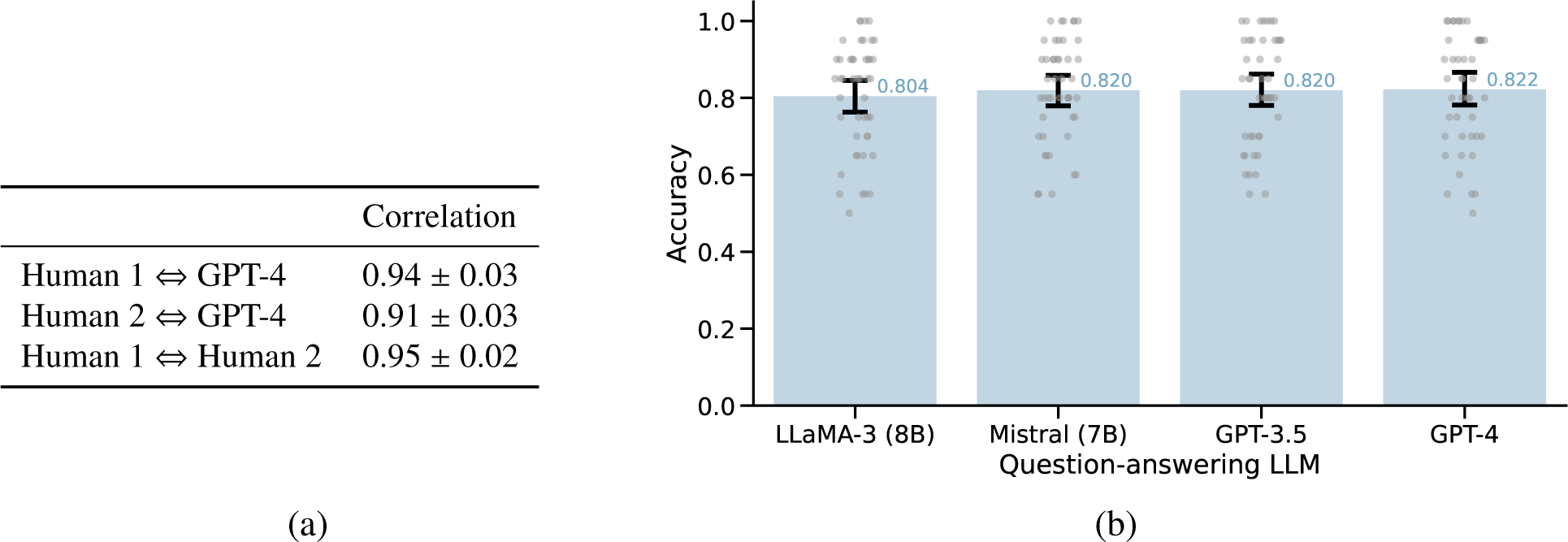
Evaluating the question-answering performance of underlying LLMs. (a) For each of the 35 stable questions, we selected 150 10-grams from the training set that had balanced answers (i.e. 100 yes answers and 100 no answers), resulting in 5,250 (question, 10-gram) pairs. We then had 2 humans manually annotate the answer for each pair. We found that, averaged across questions, the agreement (Pearson correlation) between the human annotations and GPT-4 annotations was comparable to the agreement across the human annotations. Error bars show standard error of the mean. (b) We evaluated the faithfulness of our question-answering models on a recent diverse collection of 54 binary classification datasets (*55*) (see data details in Table S3). These datasets are difficult, as they are intended to encompass a wider-ranging and more realistic list of questions than traditional NLP datasets. Each point shows an individual dataset and error bars show the 95% confidence interval. Fig. S9 shows the classification accuracy for the 3 LLMs used in our methods along with GPT-3.5 (gpt-3.5-turbo-0125). On average, each of the LLMs answered these questions with fairly high accuracy, with GPT-4 slightly outperforming the other models. However, we observe poor performance on some tasks, which we attribute to the task difficulty and the lack of task-specific prompt engineering. For example, the dataset yielding the lowest accuracy asks the question *Is the input about math research?*. While this may seem like a fairly simple question for an LLM to answer, the examples in the negative class consist of texts from other quantitative fields (e.g. chemistry) that usually contain numbers, math notation, and statistical analysis. Thus the LLMs answered *yes* to most examples and achieve accuracy near chance (50%). Note that these tasks are more difficult than the relatively simple questions we answer in the fMRI experiments, especially since the fMRI input lengths are each 10 words, whereas the input lengths for these datasets are over 50 words on average (with some inputs spanning over 1,000 words).

#### S3.5 ECoG electrode selection

**Table S3:** 54 binary classification datasets along with their underlying yes/no question and corpus statistics from a recent collection (*55*).

| Dataset name | Dataset topic | Underlying yes/no question | Examples | Unique unigrams |
| --- | --- | --- | --- | --- |
| 0-irony | sarcasm | contains irony | 590 | 3897 |
| 1-objective | unbiased | is a more objective description of what happened | 739 | 5628 |
| 2-subjective | subjective | contains subjective opinion | 757 | 5769 |
| 3-god | religious | believes in god | 164 | 1455 |
| 4-atheism | atheistic | is against religion | 172 | 1472 |
| 5-evacuate | evacuation | involves a need for people to evacuate | 2670 | 16505 |
| 6-terrorism | terrorism | describes a situation that involves terrorism | 2640 | 16608 |
| 7-crime | crime | involves crime | 2621 | 16333 |
| 8-shelter | shelter | describes a situation where people need shelter | 2620 | 16347 |
| 9-food | hunger | is related to food security | 2642 | 16276 |
| 10-infrastructure | infrastructure | is related to infrastructure | 2664 | 16548 |
| 11-regime change | regime change | describes a regime change | 2670 | 16382 |
| 12-medical | health | is related to a medical situation | 2675 | 16223 |
| 13-water | water | involves a situation where people need clean water | 2619 | 16135 |
| 14-search | rescue | involves a search/rescue situation | 2628 | 16131 |
| 15-utility | utility | expresses need for utility, energy or sanitation | 2640 | 16249 |
| 16-hillary | Hillary | is against Hillary | 224 | 1693 |
| 17-hillary | Hillary | supports hillary | 218 | 1675 |
| 18-offensive | derogatory | contains offensive content | 652 | 6109 |
| 19-offensive | toxic | insult women or immigrants | 2188 | 11839 |
| 20-pro-life | pro-life | is pro-life | 213 | 1633 |
| 21-pro-choice | abortion | supports abortion | 209 | 1593 |
| 22-physics | physics | is about physics | 10360 | 93810 |
| 23-computer science | computers | is related to computer science | 10441 | 93947 |
| 24-statistics | statistics | is about statistics | 9286 | 86874 |
| 25-math | math | is about math research | 8898 | 85118 |
| 26-grammar | ungrammatical | is ungrammatical | 834 | 2217 |
| 27-grammar | grammatical | is grammatical | 826 | 2236 |
| 28-sexist | sexist | is offensive to women | 209 | 1641 |
| 29-sexist | feminism | supports feminism | 215 | 1710 |
| 30-news | world | is about world news | 5778 | 13023 |
| 31-sports | sports news | is about sports news | 5674 | 12849 |
| 32-business | business | is related to business | 5699 | 12913 |
| 33-tech | technology | is related to technology | 5727 | 12927 |
| 34-bad | negative | contains a bad movie review | 357 | 16889 |
| 35-good | good | thinks the movie is good | 380 | 17497 |
| 36-quantity | quantity | asks for a quantity | 1901 | 5144 |
| 37-location | location | asks about a location | 1925 | 5236 |
| 38-person | person | asks about a person | 1848 | 5014 |
| 39-entity | entity | asks about an entity | 1896 | 5180 |
| 40-abbreviation | abbreviation | asks about an abbreviation | 1839 | 5045 |
| 41-defin | definition | contains a definition | 651 | 4508 |
| 42-environment | environmentalism | is against environmentalist | 124 | 1117 |
| 43-environment | environmentalism | is environmentalist | 119 | 1072 |
| 44-spam | spam | is a spam | 360 | 2470 |
| 45-fact | facts | asks for factual information | 704 | 11449 |
| 46-opinion | opinion | asks for an opinion | 719 | 11709 |
| 47-math | science | is related to math and science | 7514 | 53973 |
| 48-health | health | is related to health | 7485 | 53986 |
| 49-computer | computers | related to computer or internet | 7486 | 54256 |
| 50-sport | sports | is related to sports | 7505 | 54718 |
| 51-entertainment | entertainment | is about entertainment | 7461 | 53573 |
| 52-family | relationships | is about family and relationships | 7438 | 54680 |
| 53-politic | politics | is related to politics or government | 7410 | 53393 |

**Figure S10:**
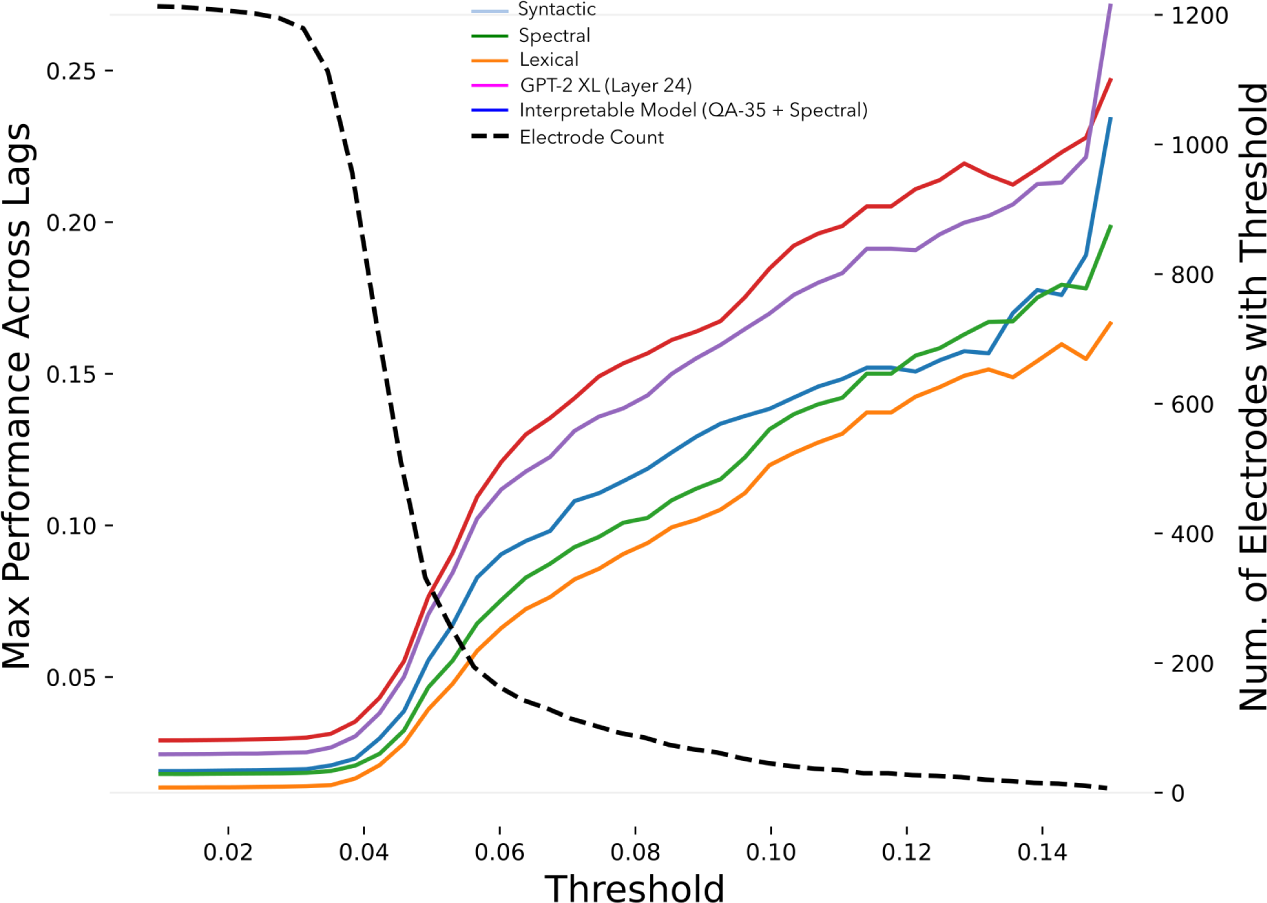
ECoG encoding model performance across test performance thresholds for selecting electrodes thresholds. The dashed line shows the number of electrodes that meet the threshold. Colored lines show the mean performance for each model from among the electrodes that meet the corresponding minimum threshold. We use a threshold of 𝑟 = 0.06 for most of our analyses, approximately at the elbow of the dashed line.

